# TLR9-activated B cells improve their regulatory function by endogenously produced catecholamines

**DOI:** 10.1101/2020.05.24.113167

**Authors:** Nadine Honke, Torsten Lowin, Birgit Opgenoorth, Namir Shaabani, Alexander Lautwein, John R. Teijaro, Matthias Schneider, Georg Pongratz

## Abstract

The sympathetic nervous system (SNS) contributes to immune balance by promoting anti-inflammatory B cells. However, whether B cells possess a self-regulating mechanism by which they modulate regulatory B cell (Breg) function is not well understood. In this study, we investigated the ability of B cells to synthesize catecholamines upon stimulation with different B cell activators. We found, that expression of the enzymes required to generate catecholamines, is upregulated by TLR9. TLR-9-specific expression of tyrosine hydroxylase (TH) correlated with upregulation of adrenergic receptors, enhanced IL-10 production, and with an overexpression of the co-inhibitory ligands PD-L1 and FasL. Moreover, concomitant stimulation of ß1-3-adrenergic receptors together with a BCR/TLR9 stimulus enhances the anti-inflammatory potential of Bregs to suppress CD4 T cells, a crucial population in the pathogenesis of autoimmune diseases, like rheumatoid arthritis. In conclusion, our data show that B cells possess autonomous mechanisms to modulate their regulatory function. These findings help to better understand the function of Bregs in autoimmune diseases and the interplay of sympathetic nervous system and B cell function.

## Introduction

The immune system requires check points to prevent reactions against self-antigens. Faulty check point inhibition can lead to multiple disorders and autoimmune diseases. While central tolerance is responsible for deleting autoreactive adaptive immune cells, peripheral tolerance is considered as a second line check point that controls lymphocytes which escaped deletion in primary lymphoid tissues. The sympathetic nervous system (SNS) might be one regulator of these processes since several studies showed that SNS influences the development and severity of autoimmune diseases, like rheumatoid arthritis (RA) ^1^. Depleting the SNS in mice showed, that the influence of the SNS is dependent on the stage of disease, with a proinflammatory role in the early phase and anti-inflammatory effect in the late phase of collagen-induced arthritis ^2^. Whereas the early proinflammatory SNS-mediated mechanisms are provided by unspecific systemic effects like increased provision of energy, higher blood and lymph flow and an increased recruitment of lymphocytes, the late anti-inflammatory effects have been proposed to be more specific on adaptive immunity with involvement of regulatory cells and IL-10 ^1,3,4^. In humans an increased activity of the SNS is observed in patients with active RA ^5^ and electric stimulation of the vagus nerve, which results among other effects in the release of sympathetic neurotransmitters in the spleen, shows some alleviating effect ^6^.

A strong neuro-immune interaction has been shown in several previous studies ^7,8^. For instance, sympathetic nerves that express tyrosine hydroxylase (TH) ^9^, the key enzyme in the biosynthesis of catecholamines, innervate lymphoid organs and gut-associated lymphoid tissues, where their nerve fiber endings are in close contact with immune cells like macrophages or T and B lymphocytes ^10,11^. While immune cell-derived cytokines regulate the sympathetic nervous system (SNS), neurotransmitters such as the catecholamine norepinephrine and neuropeptides released from the SNS can affect the function of several immune cells ^12^. Responses to catecholamines require the expression of G-protein-coupled adrenergic receptors (ADR). These adrenergic receptors are grouped into three different types (α1-, α2- and ß-ADRS) including several subtypes ^13^. The impact of these receptors on immune cells has been previously characterized ^14–16^. Among the several adrenergic receptor subtypes, the ß2-ADR was the most investigated with a bidirectional role in autoimmune diseases ^17^.

B cells are one of the major players in the pathogenesis of RA, since they can produce autoantibodies ^18,19^, and effector B cells activate CD4+ T helper cells which may worsen the situation in the affected joints ^20,21^. Regulatory B cells (Bregs) came into focus recently, since studies in mice have shown that they are not only essential to reduce chronic inflammation ^22,23^, but also effective in improving autoimmune disorders and maintaining tolerance ^3,24–26^. Several different Breg subsets have been described in mice with similarities in their surface markers and function ^27–33^. From all these Breg subclasses, B10 cells are the cell population best characterized. They are associated with IL-10 production and their crucial role in several mouse models of autoimmunity has been demonstrated ^25,26,34^. Murine B10 cells account for 1-2% of cells in the spleen and are mostly represented in the subgroup of CD19^+^CD1d^hi^CD5^+^ B cells, which comprises 10-20% of the total B cell population ^27,35^. Bregs with a phenotype of CD19^+^CD24^hi^CD27^+^ are described as one human counterpart of murine B10 cells ^36,37^. Besides IL-10 secretion, the immunosuppressive capacity or checkpoint activity of regulatory B cells has also been associated with the production of IL-35 ^38^, TGF-ß ^39^ and inhibitory costimulatory molecules e.g. PD-L1 ^40,41^ and FasL ^42,43^.

Since it is known, that the function of regulatory B cells is supported by SNS mediators, but on the other hand sympathetic nerve fibers are lost during chronic inflammation ^9,44^, we hypothesized that B cells have a self-sustained sympathetic backup mechanism to modulate their Breg function during chronic inflammation.

## Results

### B cells have their own machinery for catecholamine synthesis

Several enzymes are required to form catecholamines like norepinephrine (noradrenaline) and/or epinephrine (adrenaline) starting from the amino acid tyrosine ^45^ (Fig. 1a). The first and rate limiting enzyme in this process is tyrosine hydroxylase (TH), which converts tyrosine to L-dopa whereas phenylethanolamine N-methyltransferase (PNMT) the terminal enzyme in this cascade metabolizes norepinephrine (NE) to epinephrine (Fig. 1a). First, we tested whether B cells express the necessary enzymes for the biosynthesis of catecholamines. B cells were isolated from spleens of naïve DBA/1J mice and activated T cell-independently with anti-B cell receptor (BCR) IgM and the TLR9 agonist CpG. Interestingly, we found that naïve B cells express low levels of TH, whereas activation augmented TH levels as assessed by western blot (Fig. 1b, Supplementary Fig. 6a) and flow cytometry (Fig. 1c). Similarly, activation of B cells with anti-IgM/CpG increased protein levels of PNMT (Fig. 1d, Supplementary Fig. 6b), which converts norepinephrine to epinephrine. This demonstrates that anti-IgM/CpG-activated B cells are able to upregulate enzymes required to synthesize their own catecholamines independent from the SNS.

**Fig. 1:**
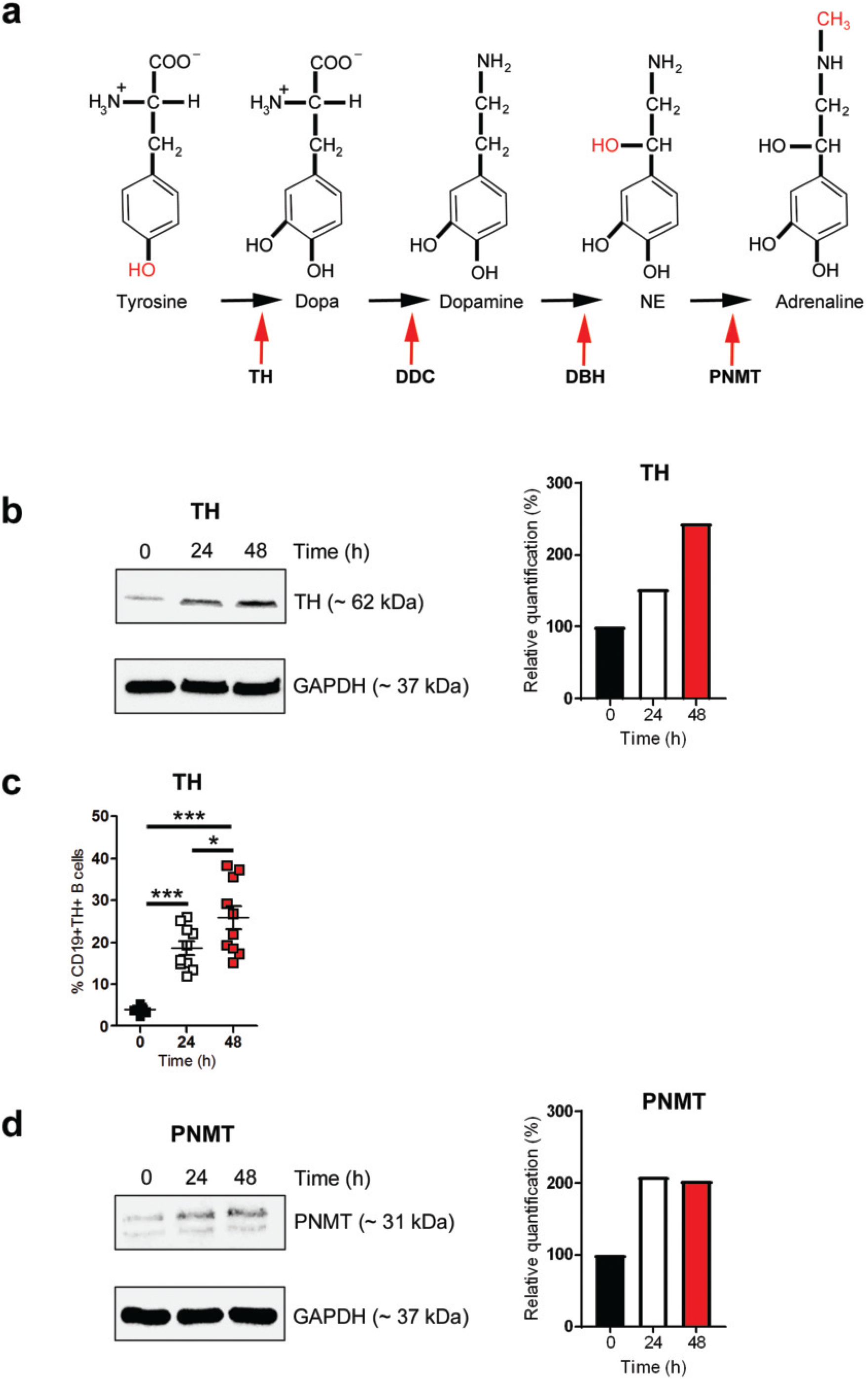
B cells have their own machinery for catecholamine synthesis. (a): Graphical depiction of the catecholamine biosynthetic pathway. (b-d): B cells were activated with anti-IgM/CpG for 24 or 48 hours or left non-activated. The expression of tyrosine hydroxylase (TH) was analyzed by western blot (b) and flow cytometry (c; n = 7-10). Expression of phenylethanolamine N-methyltransferase (PNMT) was investigated by western blot (d). One representative blot of three is shown (b, d; n = 3).The frequency of TH and PNMT expression is indicated in relation to the housekeeping protein (GAPDH; b, d). For the experiments B cells from naïve DBA/1J mice were used. Data are pooled from four experiments (c). Student’s *t*-test was used for comparisons (c). n.s. not significant; *p <0.5; ***p < 0.001.

### Catecholamines are produced in a time-/stimulus-dependent way

Catecholamines are secreted by sympathetic nerve fibers, thereby affecting the function of other cells. Some immune cells produce their own catecholamines ^46–48^, but it is yet unknown whether this is a regulated process. To investigate whether production of catecholamines depends on efficiency and persistence of activation, B cells were either activated with the BCR stimulus anti-IgM, the TLR9 agonist CpG, with both (BCR/TLR9) or were left untreated. Intracellular catecholamines were visualized by using neurosensor_521 (NS_521), a fluorescent dye that specifically reacts with dopamine and norepinephrine. After 4h of activation, B cells showed elevated levels of catecholamines compared to unstimulated B cells (Fig. 2a). Additionally, activation of B cells with anti-IgM alone resulted in minimal production of catecholamines, while most catecholamines were produced upon activation with anti-IgM/CpG (Fig. 2a). Interestingly, treating B cells with reserpine, a specific, irreversible inhibitor of the vesicular monoamine transporter 1 and 2 causes a profound depletion of endogenous catecholamines in B cells, especially during activation with CpG alone (Fig. 2a). Furthermore, treating B cells with norepinephrine (NE), an agonist acting on different α- and β-adrenergic receptors (ADRs) in a concentration-dependent way ^49,50^, reduced endogenous catecholamines in the cell, based on the activation stimulus (Fig. 2a). However, after long term activation (24h), intracellular staining showed lower levels of catecholamines, we hypothesized that catecholamines are released from the cell (Fig. 2a). In addition to flow cytometry, immunofluorescence histology supports our results that B cells upregulate intracellular catecholamines after short-term incubation whereas long-term stimulation seems to foster the release of produced catecholamines (Fig. 2b and 2c). Altogether, our results suggest that synergistic activation of B cells with anti-IgM and CpG leads to enhanced production of catecholamines.

**Fig. 2:**
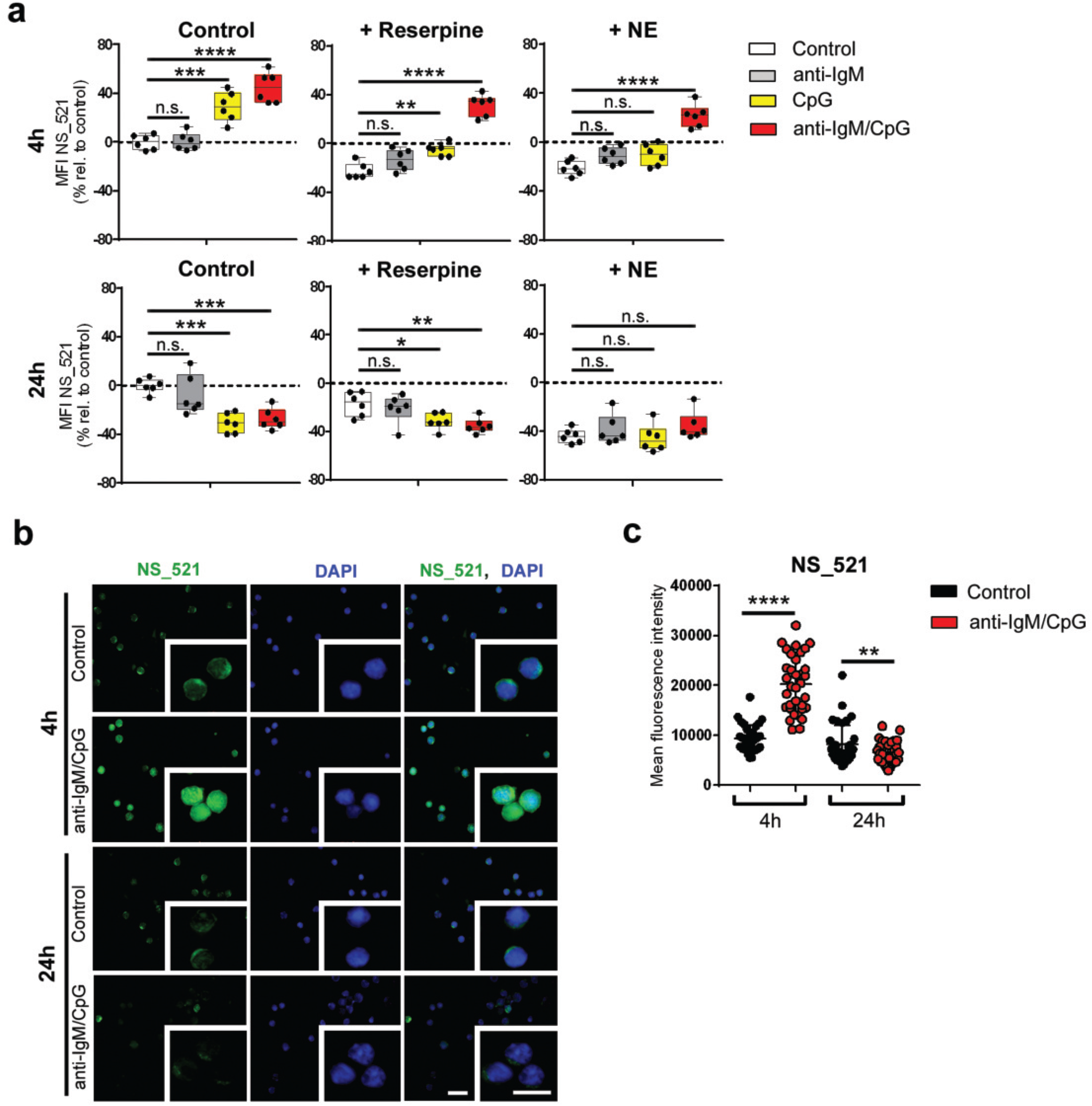
Catecholamines are produced in a time-/stimulus-dependent way. (a): B cells were activated for 4h or 24h with anti-IgM, CpG or anti-IgM/CpG and concomitantly treated with reserpine (1μM) or norepinephrine (NE; 10μM) for 1h or left untreated. As control group non-activated B cells were used. The median fluorescence intensity (MFI) of neurosensor_521 (NS_521; 5mM) was measured by flow cytometry. The frequency of MFI in relation to the control group is shown (n = 6). (b, c): Anti-IgM/CpG-activated B cells were cultivated for 4h or 24h. Non-activated B cells were used as control group. Immunofluorescence of cultivated B cells stained with NS_521 (10mM; green) and Dapi (blue). Images were captured at 40x magnification with a Zeiss AxioVision microscope. Scale bars indicate 20μm (main images) and 10μm (insets). One of three representative images is shown (n = 3). (c): The mean fluorescence intensity of B cells per condition was measured with the AxioVision software (n = 34). For the experiments B cells from naïve DBA/1J mice were used. Student`s *t*-test (c) or one-way analysis of variance (ANOVA) (a) were used for comparisons. n.s. not significant; *p <0.5; **p < 0.01; ***p < 0.001; ****p < 0.0001.

### TLR9-specific expression of tyrosine hydroxylase in B cells

It is known that B cells are activated in a T cell-dependent (TD) or –independent (TI) way by stimulation of CD40 or TLR signaling pathways. Therefore, we investigated whether upregulation of TH in B cells is also mediated by other mechanisms than TLR9 activation. For this purpose, B cells were stimulated with the TD stimulus anti-CD40 in combination with IL-4 or ligands for TLR3, TLR4 and TLR9 (TI). Unstimulated B cells served as control group. In comparison to other stimuli used, TLR9 activation induced the strongest TH expression (Fig. 3a) albeit the TLR4 ligand LPS and anti-CD40/IL-4 stimulation increased cell surface levels of MHC-II and CD86 similar to the TLR9 ligand CpG (Fig. 3b).

**Fig. 3:**
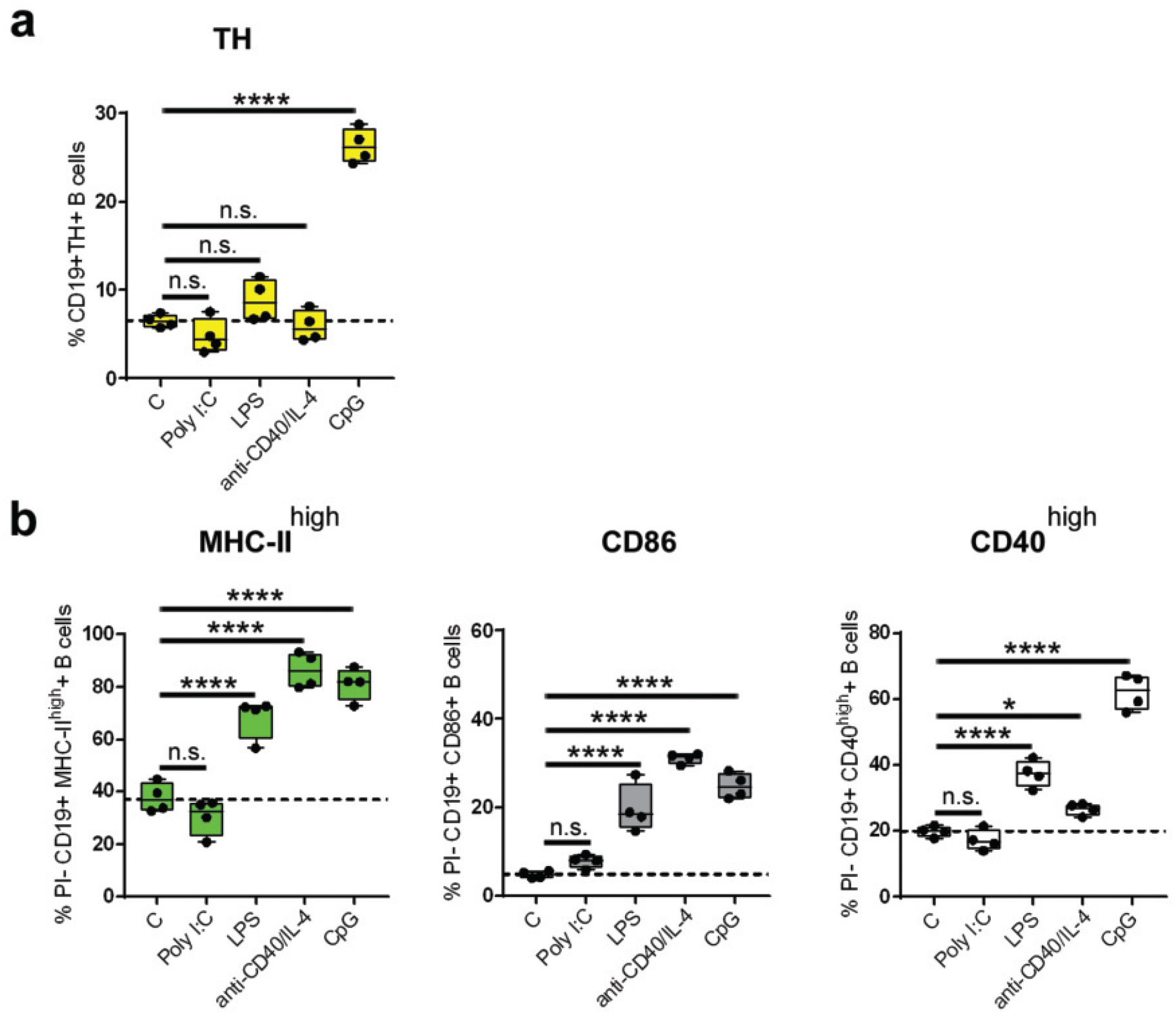
TLR9-specific expression of tyrosine hydroxylase in B cells. (a, b): B cells were activated with the T cell-dependent stimulus anti-CD40 (1,25μg/mL) in combination with IL-4 (1,25ng/mL) or with the different T cell-independent stimuli Poly I:C (TLR3; 1,25μg/mL), LPS (TLR4; 1,25μg/mL) and CpG (TLR9; 1,25μg/mL) for 24h. As control group non-activated B cells were used. The frequency of CD19+TH+ B cells (a; n = 4) and PI-CD19+MHC-II^high^+, PI-CD19+CD86+ and PI-CD19+CD40^high^+ B cells (b; n = 4) were quantified by flow cytometry. For the experiments B cells from naïve DBA/1J mice were used. One-way analysis of variance (ANOVA) (a, b) was used for comparisons. n.s. not significant; *p <0.5; ****p < 0.0001.

Interestingly, higher concentrations of TD- and TI-mitogens resulted in increased expression of activation markers (Supplementary Fig. 1a, b) and TH expression (Supplementary Fig. 2a), but none of the other stimuli induced TH as strongly as CpG (Fig. 3a). Furthermore, by testing different CpG-Oligodeoxynucleotides (ODNs), CpG-ODN 1826 was the strongest activation stimulus (Supplementary Fig. 3a) and this activation correlated with highest expression of TH (Supplementary Fig. 3b). Taken together, TH is increased in B cells specifically upon TLR9 activation.

### B cells express catecholamine receptors and transporters

Next, we wondered whether norepinephrine or epinephrine synthesized by B cells acts in an autocrine manner or is secreted to influence other cells by paracrine or endocrine mechanisms. Since the expression of adrenergic receptors is a prerequisite for autocrine stimulation with catecholamines, we determined which adrenergic receptors are expressed by B cells stimulated with the TLR9 agonist CpG. The surface expression of ADRA1A, ADRA1B, ADRA2B and ADRB2 on B cells upon activation of TLR9 agonist was determined and compared to non-activated B cells. After 48 hours, we found that TLR9-activated B cells upregulate all adrenergic receptors investigated compared to unstimulated control cells (Fig. 4a) and this was also associated with enhanced expression of TH (Fig. 4b). These data suggest that TLR9-stimulated B cells are modulated by their own monoamines in an autocrine feedback loop. Monoamine transporters control the extracellular concentration of monoamine neurotransmitters. Therefore, we tested whether B cells express catecholamine transporters and are able to regulate the uptake of catecholamines. The expression of three different transporters was investigated in B cells: the norepinephrine transporter (NET), and the vesicular monoamine transporter-1 (VMAT-1) and −2 (VMAT-2). The latter are responsible for the shuttling of catecholamines into vesicles ^51,52^ while the norepinephrine transporter (NET), mediates the reuptake of catecholamines from extracellular space ^53^. In unstimulated B cells, we detected NET on the plasma membrane, whereas VMAT-1 and VMAT2 were located intracellularly (Fig. 4c). In comparison to unstimulated B cells, activated B cells showed increased intracellular levels of all three transporters (Fig. 4d). Moreover, intracellular catecholamines were visualized by treating naïve B cells with the fluorescent dye NS_521 (Fig. 4e). The observed staining pattern suggested vesicular storage of catecholamines in B cells. The functionality of transporters was assessed by incubating non-activated and anti-IgM/CpG-activated B cells with a selective fluorescent VMAT2 substrate (FFN200 dihydrochloride) in the presence or absence of the H+-coupled VMAT blocker (reserpine). We confirmed the uptake of FFN200 by non-activated and activated B cells (Fig. 4f). Additional treatment with pan VMAT inhibitor reserpine blocked the uptake of FFN-200 resulting in reduced fluorescence intensity. In conclusion, we found that B cells have the prerequisite to respond to exogenous non-B cell-derived catecholamines via their adrenergic receptors and are able to transport and store catecholamines intracellularly for latter B cell neuron-like transmission.

**Fig. 4:**
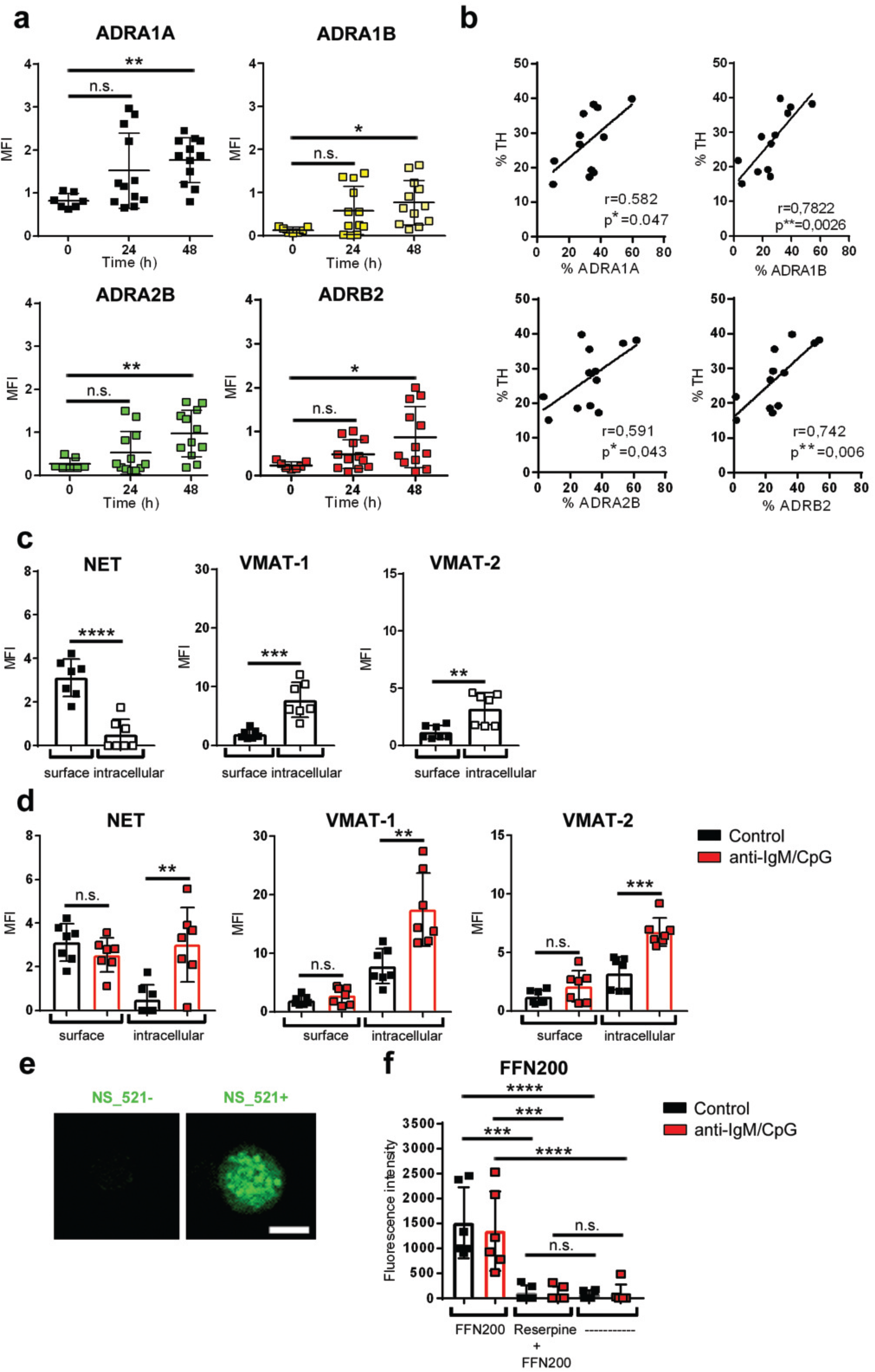
B cells express catecholamine receptors and transporters. (a, b): B cells were activated with anti-IgM/CpG for 24h and 48h. Non-activated B cells were used as control group (0h). The Median fluorescence intensity (MFI) of all adrenergic receptors (ADRs) analyzed (a; n = 7-12) was determined by flow cytometry. (b): The correlation of TH and ADR expression was quantified by flow cytometry (n = 12). (c, d): Naive B cells (c) or anti-IgM/CpG-activated (24h) B cells (d) were cultivated. The Mean fluorescence intensity (MFI) of labeled monoamine transporters was determined by flow cytometry (n = 7). (e): Non-activated B cells were stained with neurosensor_521 (NS_521; 10mM) for 30 min. at 37 ̊C and 5% CO_2_. Catecholamine producing B cells were analyzed by immunofluorescence histology. Fluorescent images were captured at 40x magnification using Zeiss AxioVision microscope. One of three representative images is shown. Scale bar = 5 μm (n = 3). (f) Non-activated and activated B cells were treated with reserpine for 45 min. or left untreated. After 30 min. the cells were incubated with FFN200 dihydrochloride (10μM) for 1.15h. FFN200 dihydrochloride- and reserpine-untreated cells were used as controls. The fluorescence intensity of FFN-200 dihydrochloride was measured by plate reader (n = 6). For the experiments B cells from naïve DBA/1J mice were used. Data are pooled from five experiments (a, b). One-way analysis of variance (ANOVA) (a, f) and Student`s *t*-test (c, d) were used for comparisons. Continuous variables were analyzed using linear regression with r values calculated by Pearson correlation (b). n.s. not significant; *p <0.5; **p < 0.01; ***p < 0.001; ****p < 0.0001.

### TH upregulation is associated with IL-10 production

In previous studies, we found ex vivo ß2-adrenergic receptor stimulation to augment B cell-derived IL-10 production in immunized mice ^4^. In this study, we confirmed that TLR9-activated B cells with anti-IgM/CpG treatment induced IL-10 expression as assessed by both qRT-PCR (Fig. 5a) and ELISA (Fig. 5b). In line with our finding that CpG increased levels of TH, we also confirmed that among different classes of CpG-ODNs, CpG1826, a class B CpG-ODN, enhanced IL-10 production (Supplementary Fig. 3c). Although higher concentrations of TD (anti-CD40/IL-4) and TI-mitogens (LPS, Poly I:C) are able to induce IL-10 production (Supplementary Fig. 2b), they could not reach IL-10 levels, observed with CpG activation (Fig. 5b). Therefore, CpG not only increases TH expression but also induces the regulatory, IL-10-producing B cell phenotype, which was also confirmed by increased percentage of CD19+CD5+CD1d+ B cells after TLR9 stimulation as determined by flow cytometry (Fig. 5c). Furthermore, enhanced expression of TGF-β and IL-35, which are markers for regulatory B cells, besides IL-10 was also demonstrated (Fig. 5d). Next, we investigated whether expression of TH is associated with IL-10 production in B cells. We employed flow cytometry and determined the frequency of CD19+TH+, CD19+IL-10+, TH+IL-10+ and IL-10+TH+ cells in non-activated and anti-IgM/CpG-activated B cells, respectively. Intriguingly, almost 80% of TH+ B cells were IL-10 positive (Fig. 5e). Moreover, inhibition of TH by using the specific tyrosine hydroxylase inhibitor 3-Iodo-L-tyrosine resulted in a significant reduction of IL-10 production by B cells (Fig. 5f) while B cell viability was not affected by this treatment (Supplementary Figure 4a, b). From these data, we conclude that TLR9 agonism specifically leads to intrinsic TH upregulation and differentiation of B cells into regulatory B cells with a concomitant increase in IL-10 production.

**Fig. 5:**
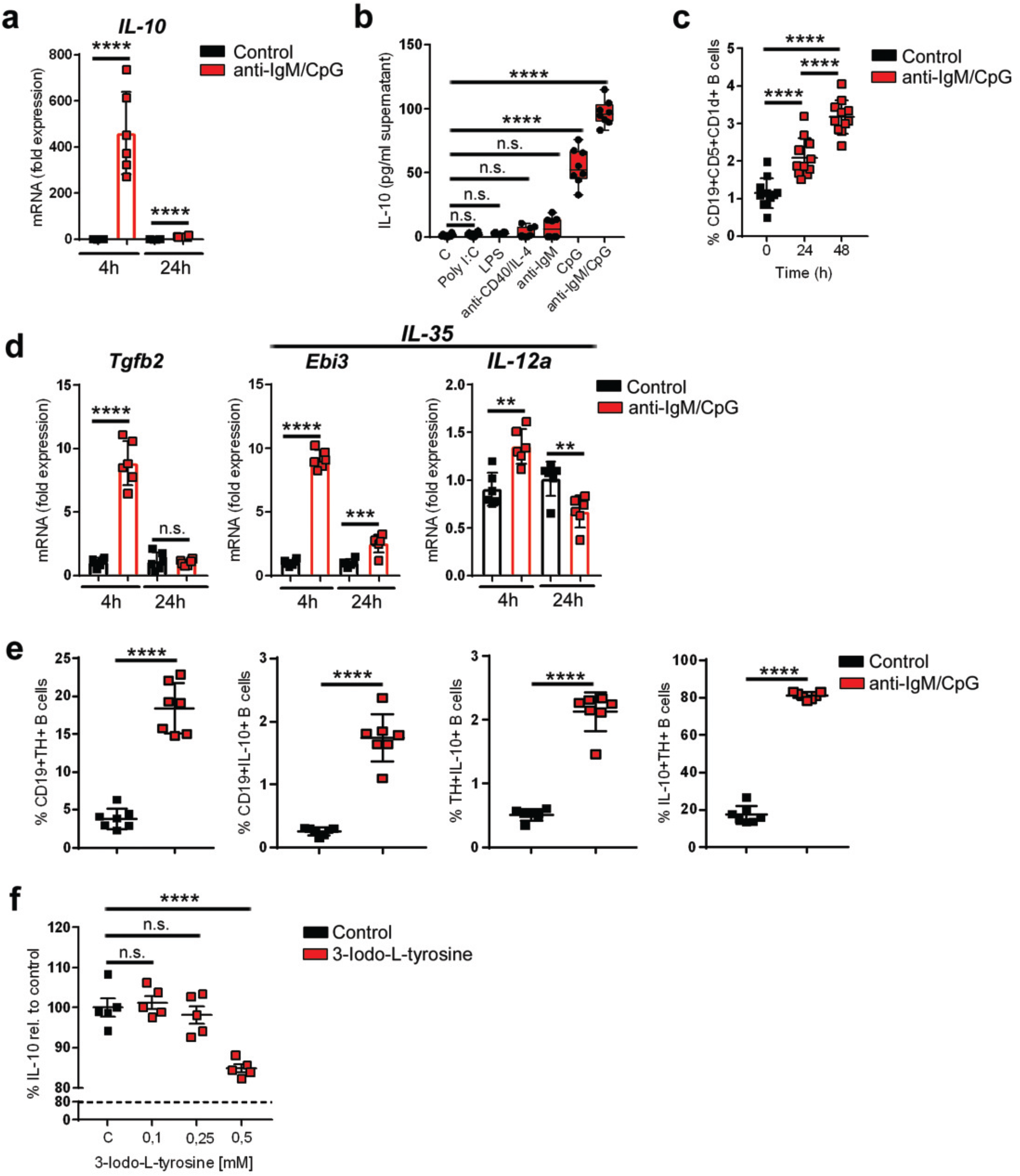
TH upregulation is associated with IL-10 production. (a): B cells were activated for 4h and 24h with anti-IgM/CpG or left non-activated. The IL-10 mRNA fold expression was determined by quantitative real-time polymerase chain reaction (qRT-PCR; n = 6). (b): B cells were activated with indicated TD- or TI-stimuli (anti-CD40; Poly I:C, LPS, CpG: 1,25μg/mL; anti-IgM: 5μg/mL; IL-4: 1,25ng/mL). Non-activated B cells were used as controls. The production of IL-10 was measured in the cell culture supernatant by ELISA (b; n = 8). (c): B cells were activated with anti-IgM/CpG or left untreated. The frequency of regulatory B cells (CD19+CD5+CD1d+) was analyzed by flow cytometry after 24h and 48h (n = 11). (d) Anti-IgM/CpG-activated B cells or non-activated B cells were cultivated for 4h or 24h. The mRNA fold expression of indicated genes was determined by qRT-PCR (n = 6). (e): B cells were activated with anti-IgM/CpG for 24h or left non-activated. The frequency of CD19+TH+, CD19+IL-10+, TH+IL-10+ and IL10+TH+ B cells was analyzed by flow cytometry (n = 7). (f): B cells were treated with different concentrations of the tyrosine hydroxylase inhibitor 3-Iodo-L-tyrosine for 30 min., before activation with anti-IgM/CpG. The production of IL-10 was analyzed in cell supernatants by ELISA (n = 5). For the experiments B cells from naïve DBA/1J mice were used. Data are pooled from three (c, e) or four (b) experiments. Student`s *t*-test (a, d, e) or one-way analysis of variance (ANOVA) was used for comparisons (b, c, f). n.s. not significant; **p < 0.01; ***p < 0.001; ****p < 0.0001.

### Activation of ß-ADRs enhance B cell-derived IL-10 production

Since anti-IgM/TLR9-activated B cells upregulate several adrenergic receptors, we checked whether adrenergic receptor signaling is important for acquiring a regulatory phenotype. For this purpose, anti-IgM/CpG-primed B cells were treated with agonists for α1-, α2-, β2- and β3-adrenergic receptors (ADRs). Activation of α1- and α2-ADRs led to a significant and concentration-dependent reduction in IL-10 production (Fig. 6a). In contrast, stimulation of B cells with subtype-specific β-ADR agonists enhanced IL-10 production (Fig. 6a). Moreover, the non-selective ß-ADR agonist isoproterenol significantly enhances IL-10 production compared to ß-subtype-specific agonists (Fig. 6b). This effect was inhibited by the pan-ß-adrenergic receptor antagonist nadolol (Fig. 6b). These data show that B cell-derived IL-10 production is dependent on catecholamine concentration, which is determined by integration of ß-ADR and α-ADR signals.

**Fig. 6:**
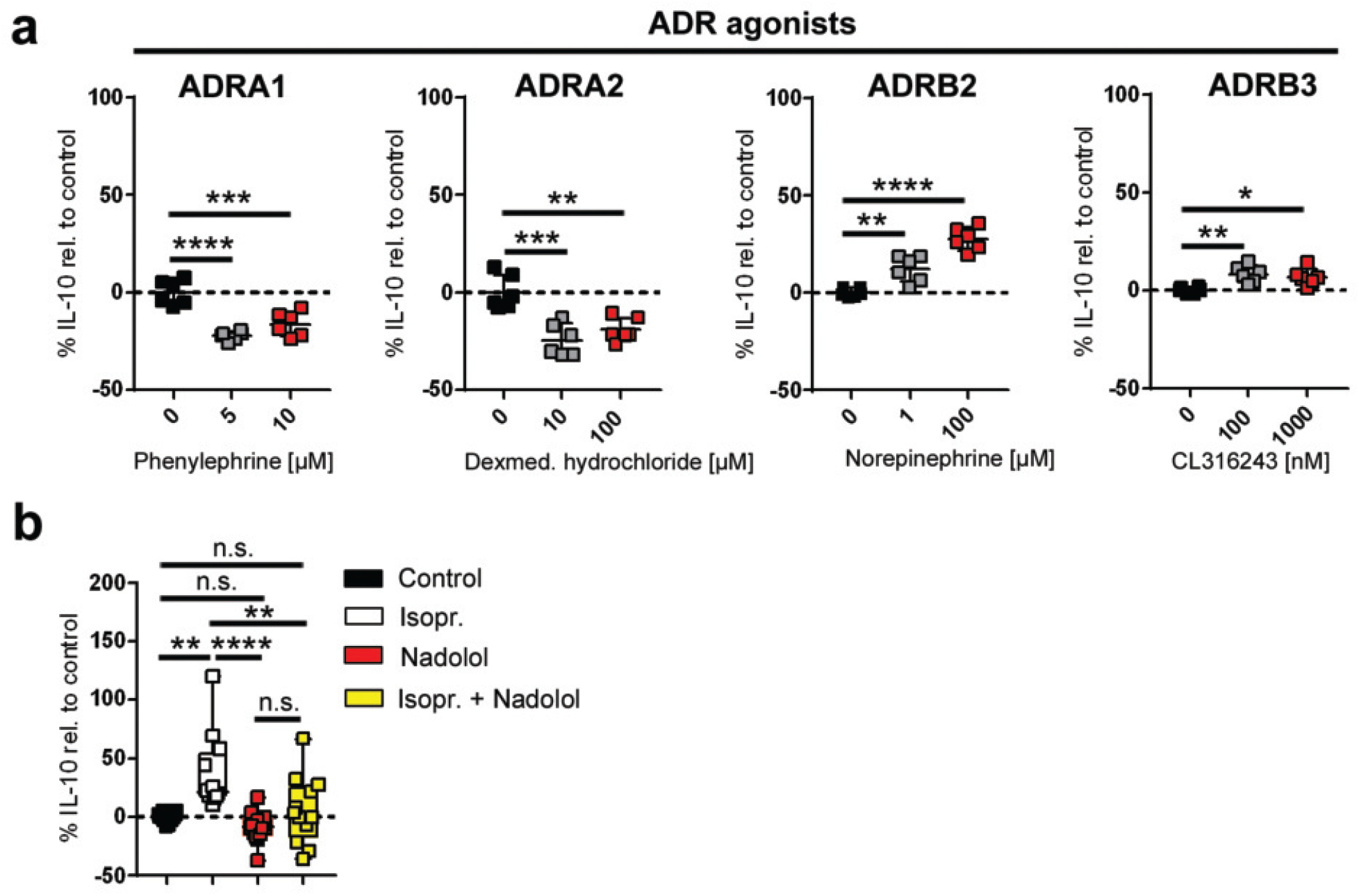
Activation of ß-ADRs enhance B cell-derived IL-10 production. (a): Anti-IgM/CpG-activated or non-activated B cells were cultivated for 4h, before additionally treated with different concentrations of α1-, α2- or β-adrenergic receptor agonists (a; n = 6). (b): B cells were either treated with the β-ADR antagonist nadolol [5μM] 30 min. prior activation with anti-IgM/CpG or were at first activated with anti-IgM/CpG for 4h before the cells were treated with the pan ß-ADR agonist isoproterenol (Isopr.; 10μM; n = 12). The production of IL-10 was analyzed by ELISA after 24h. For the experiments B cells from naïve DBA/1J mice were used. Data are pooled from five experiments (b). One-way analysis of variance (ANOVA) was used for comparisons (a, b). n.s. not significant; *p <0.5; **p < 0.01; ***p < 0.001; ****p < 0.0001.

### IL-10 improves Bregs function to suppress T cells

Regulatory B cells can inhibit or suppress inflammation mediated by CD4+ T cells, mainly T helper 1 cells (Th1) ^54,55^, which is one of the main populations that plays a crucial role in the pathogenesis of auto-inflammatory diseases like rheumatoid arthritis (RA). We wondered, whether Bregs generated by TLR9 activation with or without concomitant ß-ADR stimulation inhibit the proliferation of CD4+ T cells. TLR-9 primed B cells or naïve B cells were co-cultured with splenocytes, including activated CD4+ T cells in the presence or absence of the ß-adrenergic receptor agonists’ norepinephrine (NE) or isoproterenol (Isopr.). We found that CD4+ T cells were mainly suppressed by TLR9-induced Bregs (Fig. 7a, b). Moreover, ß-ADR stimulation of CpG-pre-activated B cells further enhanced CD4+ T cell suppression (Fig. 7a, b). Surprisingly, isoproterenol alone slightly suppressed T cell proliferation independent of CpG-activation (Fig. 7a, c). These results suggest that CpG-primed Bregs have the highest capacity to suppress inflammation induced by CD4+ T cells, which is additionally enhanced by a ß-adrenergic receptor stimulus (Fig. 7a, b). Several mechanisms are responsible for Breg-mediated immunosuppression one being the production of IL-10 and signaling via the IL-10R on target cells. To determine the role of this pathway in CpG-induced CD4+ T cell suppression, anti-IL-10 depleting antibodies were added to co-culture medium. The inhibition of IL-10 decreased the suppression of CD4+ T cells in comparison to IgG control antibodies (Fig. 7d). In addition to IL-10, Bregs impair T cell function via surface proteins such as PD-L1 ^40,41^ and FasL ^42,43^. Therefore, we determined, whether Bregs generated by TLR9 activation show enhanced expression of these proteins. Indeed, we demonstrated that both PD-L1 and FasL were significantly up-regulated in TLR9-activated B cells in comparison to untreated cells (Fig. 7e). However, this expression was not dependent on ß-adrenergic receptor stimulation but depends solely on TLR9 activation (Fig. 7e, Supplementary Fig. 5). In order to assess the impact of cell-cell interaction, cytokine production or even both in the suppression of CD4+ T cell proliferation; trans-well experiments were performed. Interestingly, in this setting, TLR9-activated B cells promoted CD4+ T cell proliferation rather than to suppress it (Fig. 7f). This increased proliferation was due to the presence of IL-10, because depletion of IL-10 significantly reduced CD4+ T cell proliferation (Fig. 7f). In conclusion, TLR9-Bregs primarily suppress CD4+ T cell proliferation in a cell-cell-contact-dependent way and that this suppression is significantly enhanced by additional presence of IL-10 and ß-adrenergic receptor signaling.

**Fig. 7:**
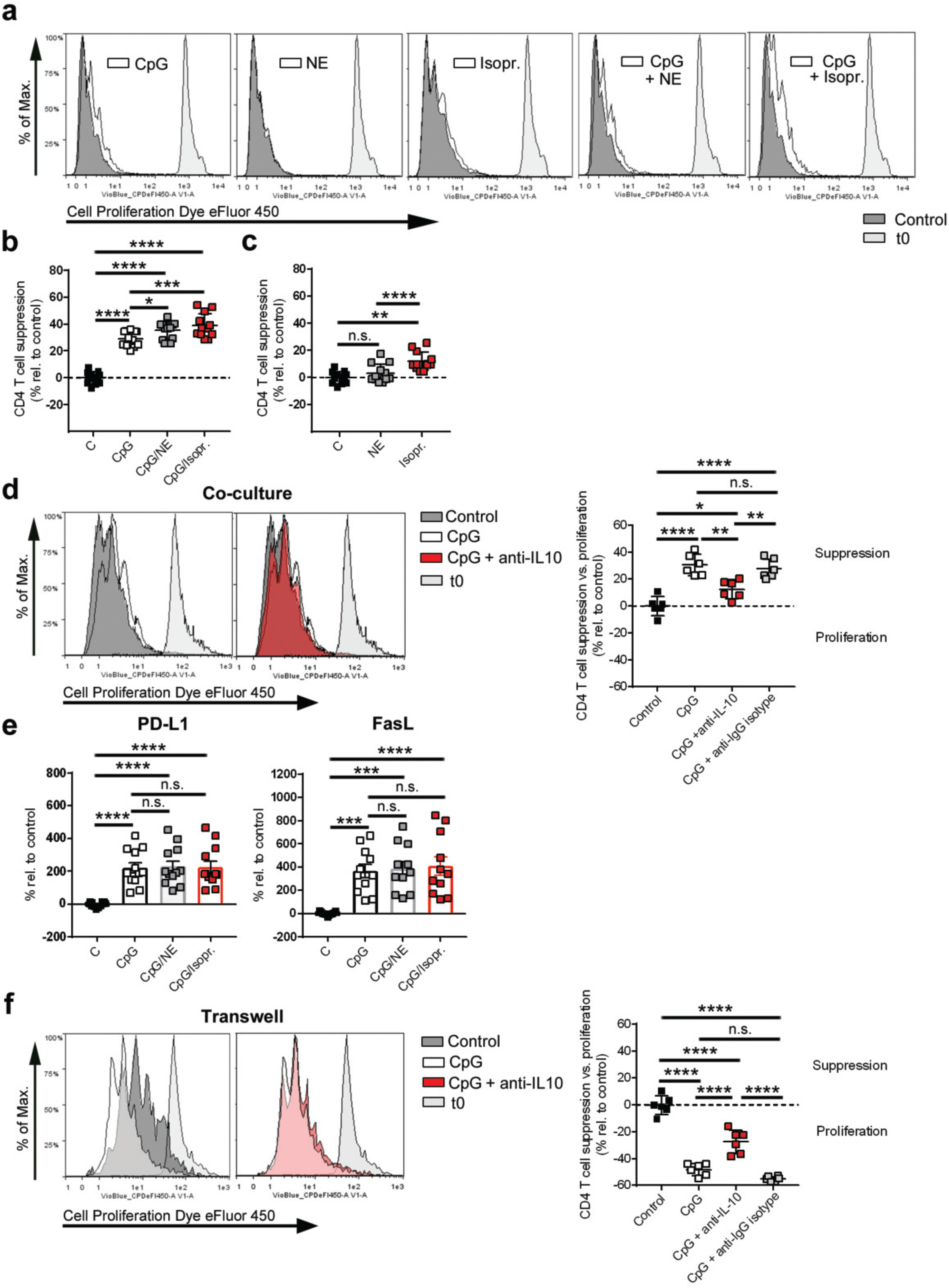
IL-10 improves Breg function to suppress T cells. (a-f): DBA/1J mice were immunized with 100μl of an emulsion of collagen type II and complete Freund`s adjuvans. After 14-21 days, splenic B cells from immunized mice were isolated by magnetic activated cell sorting (MACS) and B cell-depleted splenocytes were used for coculture and transwell experiments. (a-e): B cells were pre-activated with CpG (1,25μg/mL), isoproterenol (Isopr.; 10μM) or norepinephrine (NE; 10μM) alone or in combination before cultivation with CD3/CD28-activated splenocytes for 72h. (d): Moreover, the co-culture of CpG-pre-activated B cells and CD3/CD28-activated splenocytes was additionally incubated with or without an anti-IL-10 depletion or anti-IgG isotype control antibody. The proliferation (a, d) and suppression (b-d) of CD4+ T cells was monitored by FACS (b, c: n = 12; d: n = 6). One representative histogram is shown. The expression of PD-L1 and FasL was analyzed on co-cultured B cells after 72h by FACS (e; n = 11-12). (f): CpG-preactivated B cells and CD3/CD28-activated splenocytes were separately cultivated by using transwells and additionally treated with or without an anti-IL-10 depletion or anti-IgG isotype control antibody. After 72h the proliferation and suppression of CD4+ T cells was analyzed by FACS (n = 6). Data are pooled from four experiments (b, c, e). One-way analysis of variance (ANOVA) (b-f) was used for comparisons. n.s. not significant, *p <0.5; **p < 0.01; ***p < 0.001; ****p < 0.0001.

## Discussion

In this study, we report, that TLR9-activated B cells not only express increased levels of the catecholamine producing enzyme TH, but they actually have the ability to synthesize their own catecholamines in a time- and stimulus-dependent manner. Moreover, they employ endogenous catecholamines for autocrine signaling to influence their function and possess α- and β-adrenergic receptors as well as monoamine transporters which control uptake and storage of catecholamines. Furthermore, IL-10 production is associated and modulated by TH and the ability of TLR9-Bregs to suppress CD4+ T cells is primarily mediated by increased expression of co-inhibitory surface molecules like e.g. PD-L1 and FasL, but was additionally enhanced by elevated IL-10 production and ß-adrenergic receptor stimulation.

The interaction between the nervous system and immune system through sympathetic nerve fibers is of high importance. Lack of this interaction was shown to be involved in several immune system disorders, not only during viral infection ^56^, but also in autoimmune diseases ^57–59^. An additional backup strategy should exist to recover the lack of this interaction upon loss of peripheral sympathetic nerves during chronic inflammation ^9,44^. The ability of B cells to compensate the decreased adrenergic signaling might be considered as one of the checkpoint mechanisms to locally prevent hyperactivation of the immune system and immunopathology.

We identified that B cells are functionally able to compensate the loss of sympathetic nerve fibers in chronically inflamed tissue, since they possess proteins necessary to synthesize, sense and transport catecholamines resulting in suppression of CD4+ T cell proliferation. Whether B cells alone take over catecholaminergic signaling from lost sympathetic nerves in chronically inflamed joints and tissues or need the support of other cells has to be investigated in further studies. In this context, questions arise about the amount of catecholamines produced by B cells and about the concentration needed to fully compensate the loss of sympathetic nerve fibers. However, we show that blocking TH results in decreased IL-10 production in a pure B cell culture underscoring the importance of B cell TH activity self-sustained B cell catecholamine production.

In this study, we employed neurosensor_521 (NS_521), to directly detect and monitor catecholamine content in B cells. With this fluorescent dye, we demonstrated for the first time, that catecholamines are detectable in B cells by flow cytometry and that they are stored in synaptic-like vesicles by a VMAT-mediated uptake mechanism thus mimicking further functional aspects of neuronal tissue. Monitoring of catecholamine content was previously only performed in chromaffin cells and only detectable with methods like HPLC ^60,61^ or capillary electrophoresis assays with electrochemical detection ^46,47^. Therefore, this is the first time to directly observe catecholamine metabolism in immune cells.

Adrenergic receptors are also expressed on other immune cells such as T cells ^62^. Different studies showed that signaling through these receptors is important to regulate their function and an impaired signaling of β2-AR in circulating T cells promotes immunological diseases such as RA ^15,17,63^. It remains to be determined whether T cells benefits from catecholamines produced from B cells to show a modulation into a regulatory phenotype. Some authors have proposed that also T cells express TH ^64,65^, and similar mechanisms as described for B cells in the present report, might also be valid for T cell or other TH expressing immune cells. However, there are also immune cells, like macrophages, that do express transporters for catecholamines, but are not capable of synthesizing catecholamines on their own ^66^. This argues for a differentiated and independent sympathetic system throughout the immune system, which is not necessarily controlled by upper level systems like the CNS.

Bregs influence various immune cells, for example, by inhibiting the differentiation of Th1 and Th17 cells ^54,55^, or by affecting cytokine production of monocytes and dendritic cells ^54,67^, while promoting the expansion of regulatory T cells ^68–70^. In coculture experiments, we found that TLR9-activated Bregs are able to inhibit the proliferation of CD3/CD28-activated CD4+ T cells via the expression of inhibitory cell-cell-contact molecules. The strongly upregulation of PD-L1 and FasL expression on B cells after TLR9 activation suggests that these inhibitory molecules are crucial to inhibit CD4+ T cell proliferation. Furthermore, we were able to demonstrate that this suppression effect by cell surface molecules was further enhanced by the production of IL-10 and ß-adrenergic receptor signaling. Separation of activated B cells and splenocytes through a membrane (transwell), did not lead to suppression of CD4 T cells, but *vice versa* increased T cell proliferation. This indicates that soluble factors such as IL-10 are not self-sufficient to directly augment T cell suppression but enhance cell surface signals. This hypothesis is supported by a previous study demonstrating that treatment of activated T cells with different concentrations of recombinant IL-10 does not directly inhibit proliferation, but first affects antigen-presenting cells (APCs), which then suppress T cells ^71^. Moreover, the modulating effect of IL-10 on B cells was already demonstrated in earlier studies ^72,73^, whether IL-10 influences directly the expression of cell inhibitory molecules e.g. PD-L1, FasL, to increase cell contact-dependent suppressive activity of B cells ^41,43,74–76^, is currently unknown but suggested by previous publications showing an IL-10-inducible PD-L1 expression on monocytes ^77,78^. Another possibility is that IL-10 is able to directly affect T cell suppression, but only in the presence of signaling from cell inhibitory molecules; however, this needs further investigation.

In several experimental mouse models of autoimmune diseases, a reduced regulatory B cell function has been observed ^34^. This is in line with studies in humans, showing a reduction or functional impairment of IL-10-producing Bregs in active autoimmune diseases like systemic sclerosis ^79,80^, ANCA-associated vasculitis ^81,82^, rheumatoid arthritis (RA) ^83–85^ or systemic lupus erythematosus ^75^. Observations show that the Breg numbers return to normal levels when patients are in remission. From these data, it is suggested that augmenting Breg function or number *ex vivo* and transferring them back into the host ameliorates disease and restores the balance between proinflammatory and anti-inflammatory signals. Indeed, in some experimental mouse models it has been validated that transfer of Bregs back into the host improves several autoimmune diseases including collagen-induced arthritis (CIA) ^34,86^, experimental autoimmune encephalomyelitis (EAE) ^87^ or lupus ^30^. A positive role of the SNS in this context is supported by previous experiments showing that only transfer of splenic B cells from mice with an intact sympathetic nervous system and not B cells from sympathectomized mice ameliorates collagen-induced arthritis ^4^.

In the present study, we revealed that more than 80% of IL-10-producing B cells also express TH and only less than 20% of TH+ cells do not produce IL-10, recommending TH expression as a new biomarker for regulatory B cells. A previous study confirmed that there is a general connection between TH expression and the anti-inflammatory properties of cells, since adoptive transfer of *in-vitro* generated TH +-dependent anti-inflammatory neuronal cells leads to significant reduction of CIA in mice ^88^, however, the exact mechanism by which TH expression leads to amelioration of disease was not reported, but it is hypothesized that catecholamine production is important. These findings are in line with our results, since activated B cells increase TH, produce catecholamines and augment suppressive action via increased IL10. However, our results also show that the effect of catecholamines on B cell function depends on the pattern of adrenergic receptor subtypes expressed and therefore is not static but might depend on context factors that determine these patterns. While additional stimulation via ß-adrenergic receptors significantly improved regulatory B cell function, activation via α-adrenergic receptors led to reduced amount of IL-10 produced. The fact that ß-ADR signaling pathways play an important role in increasing IL-10 was supported by other recent publications ^89,90^. In this context, we were able to show that simultaneous activation of all ß-ADR subtypes, not only β2-ADR, synergistically augment the amount of IL-10 produced. These results indicate that there is a strong link between the amount of catecholamines and the anti-inflammatory potential of B cells. The findings may be helpful in the development of new therapeutic strategies, like the *in vitro* generation of Bregs and subsequent autologous transfer or by including ß-adrenergic receptor agonists or α-adrenergic receptor antagonists in current therapeutic regimens to increase regulatory B cell function in autoimmunity. On the other hand, in case of infection, inhibiting TLR9 signaling on B cells and/or blocking β-ADRs is useful to enhance immune responses.

In this study, we showed for the first time, that TLR9-activated B cells differentiate into catecholamine producing, sympathetic neuron-like units that synthesize, store, take up and respond to catecholamines, which are important for modulating Breg function in an autocrine and/or paracrine manner by functional ß- and α-ADRs. Furthermore, we contributed to the general understanding of Breg function by presenting data that show that IL-10 is not self-sufficient to promote immunosuppression and rather augments cell-contact dependent inhibitory mechanisms and TH might be considered as additional marker for murine regulatory B cells. Taken together, this intriguing, self-sustained catecholamine system in B cells but also other immune cells might be exploited to develop new treatment strategies for immune-mediated diseases.

## Materials and methods

### Ethics Statement

Animal experiments were carried out with approval of the “Landesamt für Natur, Umwelt und Verbraucherschutz Nordrhein-Westfalen” (LANUV), Germany (approval number: 84-02.04.2016.A283) in accordance with the German laws for animal protection. Animal care and documentation was supervised by the “Zentrale Einrichtung für Tierforschung und wissenschaftliche Tierschutzaufgaben (ZETT)” of the Heinrich-Heine-University Düsseldorf, Düsseldorf, Germany.

### Mice

All experiments were performed with animals housed in single ventilated cages, under the authorization of the Veterinäramt Nordrhein Westfalen (Düsseldorf, Germany) and in accordance with the German law for animal protection. Male DBA/1J mice, 6-10 weeks old were originally purchased from Janvier Labs (Roubaix, France). Animals were housed 5 in each cage and were fed standard laboratory chow and water ad libitum under standard conditions of temperature and light. Animals care was in accordance with institutional guidelines. All experimental protocols were approved by the Landesamt für Natur, Umwelt und Verbraucherschutz (LANUV) Nordrhein-Westfalen.

### Collagen-induced arthritis

Male DBA/1J mice (8-10 weeks old) were intradermally immunized at the base of their tails with 100μl emulsion containing bovine type II collagen (2mg/ml, Chondrex, Redmond, WA, USA) emulsified in an equal volume of complete Freund`s adjuvant (CFA; 2mg/mL, Chondrex) to generate collagen-induced arthritis. Preparation of the emulsion was done according to the manufacturer`s protocol. Mice were used 14-21 days after the initial injection.

### Antibodies

**Primary antibodies**: anti-Alpha-1A adrenergic receptor (ADRA1A, Abcam, Cambridge, UK, Clone: EPR9691(B); anti-Alpha-1B adrenergic receptor (ADRA1B, Abcam, Clone: EPR 10336); anti-Alpha-2B adrenergic receptor (ADRA2B, Abcam, Clone: EPR9623); anti-Beta-2 adrenergic receptor (ADRB2, Abcam, Clone: EPR707(N)); anti-CD4-APC-Vio770 (Miltenyi Biotec, Bergisch Gladbach, Germany, Clone: REA604); anti-CD19-PE (Miltenyi Biotec, Clone: 6D5); anti-CD19-VioBright-Fitc (Miltenyi Biotec, Clone: 6D5); anti-CD40-APC (Miltenyi Biotec, Clone: FGK45.5); anti-CD86-APC-Vio770 (Miltenyi Biotec, Clone: PO3.3); anti-Fas Ligand-APC (FasL, CD178, eBioscience TM, ThermoFisher Scientific, Clone: MFL3); anti-Interleukin-10 (IL-10-PE, Miltenyi-Biotec, Clone: JES5-16E3); anti-Interleukin-10 (R&D Systems, Minneapolis, Minnesota, USA; Clone: JES052A5) for depletion; anti-Major Histocompatibility Complex II (MHC-II-PE (Miltenyi Biotec, Clone: REA610); anti-Norepinephrine transporter (NET, Alomone, catalogue number: AMT-002); anti-Programmed death ligand 1 (PD-L1-PE/Cy7, BioLegend, London, United Kingdom, Clone: 10F.9G2); anti-Tyrosine hydroxylase (TH, FACS: Abcam, Clone: EP1533Y; anti-Vesicular monoamine transporter-1 (VMAT-1, Alomone, catalogue number: AMT-007); anti-Vesicular monoamine transporter-2 (VMAT-2, Alomone; catalogue number: AMT-006). **Secondary antibody**: goat anti-rabbit IgG H&L-Alexa Fluor 488 (Abcam, catalogue number: ab150077). **Isotype controls**: rabbit IgG-PE, monoclonal (Abcam, Clone: EPR25A); rat IgG2b-APC-Vio770 (Miltenyi Biotec, Clone: ES26-5E12.4); rabbit IgG, monoclonal (Abcam, Clone: EPR25A); rabbit IgG, polyclonal (Abcam, catalogue number: ab37415); REA control-APC (Miltenyi Biotec, Clone: REA293); REA control-PE (Miltenyi Biotec, Clone: REA293); rat IgG kappa (eBioscience™, ThermoFisher Scientific, Clone:eBRG1; rat IgG 2b k-PE/Cy7 (Biolegend, Clone: RTK4530); rat IgG 2b k-PE (Miltenyi Biotec, Clone: ES26-5E12.4); armenian hamster IgG-APC (eBioscience™, ThermoFisher Scientific, Clone: eBio299Arm); rat IgG2a-PE (Miltenyi Biotec, Clone: ES26-15B7.3); rat IgG2a-APC (Miltenyi Biotec, Clone: ES26-15B7.3); rat IgG2a-VioBright-Fitc (Miltenyi Biotec, Clone: ES26-15B7.3). REA control-APC-Vio770 (Miltenyi Biotec, Clone: REA293).

### B cell isolation

Splenic mouse B cells from naïve or immunized DBA/1J mice were isolated by magnetic-activated cell sorting (MACS) via negative selection with the Pan B cell isolation Kit (Miltenyi Biotech, Bergisch Gladbach, Germany) according to the manufacturer`s protocol. The purity was controlled by flow cytometry and was 98%. B cell-depleted splenocytes were used for T cell suppression assay.

### Cultivation of B cells

Splenic mouse B lymphocytes were cultivated in Roswell Park Memorial Institute (RPMI) 1640 medium (Gibco, ThermoFisher Scientific, Paisley, United Kingdom) containing GlutaMax (200mM L-alanyl-L-glutamine-dipeptide), 25mM HEPES (2(−4-(2-hydroxyethyl)-1-piperanzinyl) ethansulfonacid) and supplemented with 10% (v/v) fetal bovine serum (FBS; Gibco, ThermoFisher Scientific), 10.000 U/mL (v/v) Penicillin-Streptomycin (Gibco, ThermoFisher Scientific) and 0,1 % (v/v) 2-mercaptoethanol (ß-ME, Merck, Darmstadt, Germany). B cell cultures were maintained by 37°C containing 5% CO_2_.

### Activation of B cells

B cells were treated with different concentrations of T cell-independent (TI) mitogens (Lipopolysaccharide (LPS; Sigma-Aldrich, Steinheim, Germany catalogue number: L2630), Polyinosinic-polycytidylic acid sodium salt (Poly I:C, Sigma, catalogue number: P1530), CpGs (InvivoGen, tlrl-kit9m) or T cell-dependent (TD) mitogens (anti-CD40 (Invitrogen, Carlsbad, CA, USA; Clone: 1C10) in combination with or without recombinant interleukin-4 (IL-4, Miltenyi Biotec, catalogue number: 130-094-061) for 24h. For B cell receptor (BCR) activation, cells were treated with 5μg/mL anti-IgM F(ab`)2(μ chain) fragment (Jackson ImmunoResearch, West Grove, PA, catalogue number: 115-006-075).

### Treatment of B cells with different ADR agonists

For the *in vitro* stimulation with adrenergic receptor ligands, splenic B cells were seeded at 2,5×10^5^ cells per well in a 96-well plate. B cells were either incubated with the ß-ADR antagonist nadolol [5μM, Sigma Aldrich] 30 min prior to activation followed by treatment with 5μg/mL anti-IgM and 1,25μg/mL CpG or were first activated with anti-IgM/CpG for 4h followed by incubation with indicated concentrations of α- or ß-adrenergic receptor (ADR) agonists (ADRA1: (R)-(-)-Phenylephrine hydrochloride (Tocris, Bristol, UK, catalogue number: 2838), ADRA2: Dexmedetomidine hydrochloride (Tocris, catalogue number: 2749), ADRB2: L-(-)—Norepinephrine (+)-biartrate salt monohydrate (Sigma-Aldrich, catalogue number: A9512), ADRB3: CL316243 disodium salt (Tocris, catalogue number: 1499), ADRB: (-)-Isoproterenol hydrochloride (Sigma-Aldrich, catalogue number: I6504). IL-10 production was analyzed in the supernatant after 24h.

### Functionality of monoamine transporter VMAT-1/2

To test the functionality of the catecholamine transporters VMAT-1/2, anti-IgM/CpG-activated or non-activated B cells were incubated with or without reserpine (5 μM, Tocris, catalogue number: 2747), an inhibitor of VMAT-1 & VMAT-2 for 45min. Then, B cells were incubated with FFN200 (10μM, Tocris, catalogue number: 5911), a selective substrate for the vesicular monoamine transporter-2 for 1.15h. Fluorescence intensity was quantified in the multimode plate reader Infinite 200 PRO (Tecan, Männedorf, Switzerland).

### Total RNA extraction, cDNA synthesis, and quantitative real-time PCR

RNA was isolated from MACS-sorted splenic B cells with the RNA Mini Kit (Qiagen, Hilden, Germany). Quantification of RNA was performed with a NanoPhotometer (Implen, Munich, Germany). The RNA was reverse transcribed to cDNA with the Quantitect Reverse Transcription kit (Qiagen). Gene expression analysis was performed with primer assays from Qiagen [*IL-10*] and Eurofins [*IL-12α, Ebi3; Tgfb2, B2m*; table 1]. For quantitative analysis, the expression levels of all target genes were normalized against beta-2-microglobulin (*B2m*; Δ cycle threshold [Ct]). For *IL-10*, *IL-12α*, *Ebi3* and *Tgfb2* gene expression values were then calculated by the delta delta Ct (ΔΔCt) method, with the mean of the control group being the calibrator to which all other samples were compared. Relative quantities (RQs) were determined with the equation RQ = 2^−ΔΔCt^.

**Table 1:**
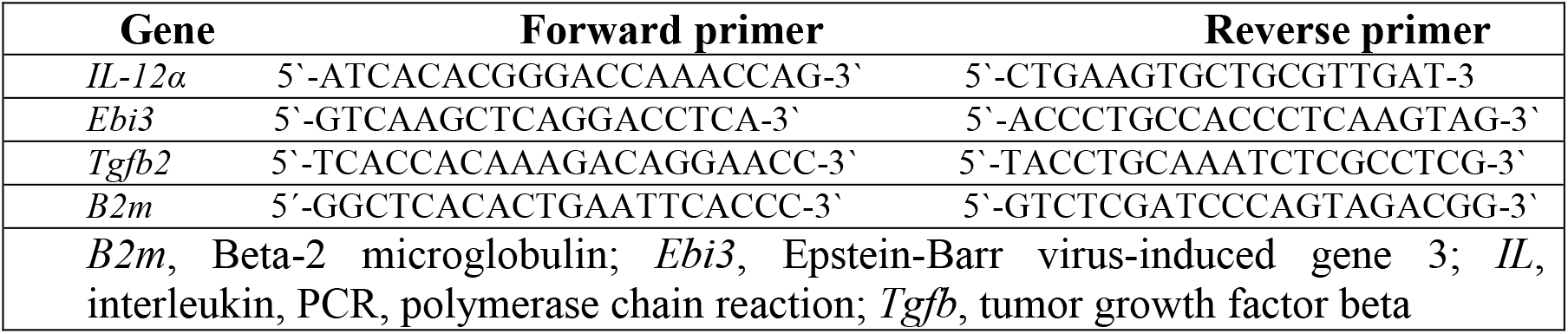
Primers used for quantified reverse transcription PCR.

### Immunofluorescence histology

For the visualization of catecholamines, B cells were treated with anti-IgM/CpG for 4h and 24h or left untreated. Afterwards cells were fixed in 2% PBS/formaldehyde before staining with neurosensor_521 (NS_521; 10mM, green) for 30 min at 37°C. The nucleus was stained with DAPI (blue) which was supplemented in ProLong Gold reagent (Invitrogen, Carlsbad, Germany).

### Inhibition of tyrosine hydroxylase (TH)

Tyrosine hydroxylase activity was inhibited by treating B cells with different concentrations of 3-Iodo-L-tyrosine (Sigma-Aldrich, catalogue number: 18250), 30 min before activation with anti-IgM/CpG.

### Flow cytometry

Splenic B cells were fixed with 2% PBS/formaldehyde for 20 min. at room temperature (RT) and treated with FcR blocking reagent (Miltenyi Biotec, catalogue number: 130-092-575) for 10 min. to inhibit unspecific Fc receptor bindings. For cell surface staining, B cells were either incubated with the primary antibodies anti-NET, anti-VMAT-1 and anti-VMAT-2 to detect catecholamine transporters or with anti-ADRA1A, anti-ADRA1B, anti-ADRA2B, and anti-ADRB2 to evaluate the expression of adrenergic receptors. After 30 min. cells were incubated with the secondary antibody Alexa488-conjugated goat anti-rabbit (1:2000) for 30 min. at 4°C. In addition, PE-conjugated anti-CD19 antibody was added for 10 min.

For intracellular staining of catecholamine transporters and tyrosine hydroxylase (TH), B cells were additionally permeabilized with Inside Perm (Miltenyi Biotec) and stained with the unlabeled primary antibodies anti-NET, anti-VMAT-1 and anti-VMAT-2 or with the directly anti–TH-PE-conjugated antibody. After 30 min., cells were either incubated with the secondary goat anti-rabbit Alexa488-labeled antibody (1:2000, for catecholamine transporters) for 30 min. or were directly analyzed (TH) with the MacsQuant® Analyzer 10 (Miltenyi Biotec). For intracellular cytokine staining B cells were activated or non-activated with anti-IgM/CpG in the presence of 5 mg/mL brefeldin A (BFA, Sigma, catalogue number: B6542) to prevent cytokine release. After 12 h B cells were incubated with anti-CD19-VioBright FITC antibody for 10 min. at 4°C, fixed with 2% PBS/formaldehyde for 20 min. at RT and permeabilized with InsidePerm at 4°C (Miltenyi Biotec) followed by incubation with anti-IL-10-PE-conjugated antibody (Miltenyi Biotec). For all primary antibodies relevant isotype control antibodies were used. Stained B cells were investigated with the MACSQuant Analyzer 10 (Miltenyi Biotec) and analyzed by using Flow-Jo or Flowlogic software.

For the detection of activation markers, unfixed B cells were incubated with anti-CD86-APC-Vio770, anti-MHC-II-PE and anti-CD40-APC in combination with anti-CD19-VioBright FITC for 10 min.

### IL-10 ELISA

Enzyme-linked immunosorbent assay (ELISA; BD OptEIA™, San Diego, CA, USA) was conducted according to manufacturer`s instructions.

### T cell suppression assay

Splenocytes from immunized DBA/1J mice were labeled with the cell proliferation dye eFluor 450 (10μM; eBioscience™, ThermoFisher Scientific, catalogue number: 65-0842-90) and activated with 1μg/mL soluble anti-CD3e (eBioscience™, ThermoFisher Scientific, Clone: eBio500A2) and 1μg/mL soluble anti-CD28 (eBioscience™, ThermoFisher Scientific, Clone: 37.51) antibody. Autologous B cells were preactivated with CpG (1,25 μg/mL) for 4h or left non-activated in the presence or absence of ß-adrenergic receptor agonists norepinephrine (10μM) or isoproterenol (10μM). After pre-activation, B cells were washed twice with PBS before using them for co-culture and transwell experiments. Activated and eFluor450-labeled splenocytes were co-cultured in a 96 well plate (Greiner Bio-One, Solingen, Germany) with or without preactivated B cells at ratio of 1:2 in cRPMI medium, for 72h at 37°C and 5 % CO_2_. For testing the influence of IL-10 and the expression of PD-L1 and FasL in the suppression of CD4+ T cells, B cells were either cocultured with splenocytes or separated by them by performing transwell experiments. For transwell experiments, pre-activated B cells were placed into the upper chamber of a 96-well insert and the activated, eFluor450-labled splenocytes were added into the lower chamber, separated with a membrane (pore size: 3μm, Sigma-Aldrich) and both containing cRPMI medium. To evaluate the direct effect of IL-10 on proliferation of CD4+ T cells an anti-IL-10 depletion antibody (5μg/mL) and rat IgG kappa (5μg/mL) isotype control antibody was used in co-culture and transwell experiments. After 72h cells were incubated with anti-CD19-VioBright-Fitc, anti-PDL1-PE/Cy7 and anti-FasL-APC to analyze surface markers on B cells involved in T cell suppression and with CD4-APC-Vio770 to investigate T cell proliferation by flow cytometry (MACSQuant Analyzer 10, Miltenyi Biotec).

### Annexin V-FITC Binding Assay

Untreated (control) and 3-Iodo-L-tyrosine-treated B cells were labeled by using the Annexin V Fitc Apoptosis detection kit (BD, Heidelberg, Germany) according to manufacturer`s instructions. Before analysis propodium jodide (PI, Miltenyi Biotec) was added to B cells and then analyzed by flow cytometry (MacsQuant Analyzer 10; Miltenyi Biotec,). The double labeling procedure allows a distinction into living (Annexin V −/PI−), early apoptotic (Annexin V +/PI−), late apoptotic (Annexin V+/PI+) and necrotic cells (Annexin V −/PI+).

### Western Blot analysis

Isolated splenic B cells from naïve DBA1/J mice, were homogenized in lysis buffer (50 mM Tris-HCl, pH 8.0; 150mM NaCl; 2mM EDTA; 1% NP-40; 0,1% SDS; 1% sodiumdeoxycholate; 1mM sodiumorthovanadate; 1 mM PMSF; 50mM sodiumfluoride; protease-inhibitor-cocktail (1x)) for 5 min on ice, snap-frozen in liquid nitrogen (freeze/thaw cycle 2x) and centrifuged for 10 min. by 10.000 g at 4°C. Protein concentration was determined in the cell supernatant, mixed with Rotiload (1:4, BIO-RAD, Feldkirchen, Germany) and boiled for 5 min. at 95°C. Total B cell protein (10μg) was separated by 12,5% SDS polyacrylamide gel electrophoresis (SDS-PAGE) and blotted at 30 mA for 90 min. on nitrocellulose membranes (BIO-RAD). After blocking with 5% (w/v) blotting-grade nonfat dry milk (BIO-RAD) for 1h at RT, membrane was washed with TBST, and incubated overnight at 4°C followed by incubation with primary antibody: anti-tyrosine hydroxylase (TH, 1:1000, Merck Millipore, Darmstadt, Germany, AB152). After washing with TBST, membrane was incubated with horseradish peroxidase (HRP)-conjugated secondary antibody (1:2000, Dako, Hamburg, Germany, catalogue number: P0448) for 2h at RT. As reference, GAPDH was used to calculate the levels of target proteins. For the visualization the proteins ECL-solution (Luminol, Para-Hydroxycoumarinacid, DMSO) and ChemiDoc Touch™ Imaging System (BIO-RAD) was used. Protein spots were analyzed with the Image Lab 5.2.1 software (BIO-RAD).

### Statistics

If not otherwise stated, data are represented as mean ± S.D. Unpaired two-tailed student’s *t*-test and one-way analysis of variance (ANOVA) was applied using GraphPad Prism software to calculate statistically significant differences between groups. The level of statistical significance was set at n.s. not significant, \**p* < 0.05, \*\**p*< 0.01, \*\*\**p*< 0.001 or \*\*\*\**p* < 0,0001. Each data point reflects results from B cells from one mouse. Data are pooled from at least two experiments unless mentioned otherwise in the figure legend.

## Supporting information

Increased concentrations of TD-/TI-stimuli raise B cell activation

Increased concentrations of TD-/TI-stimuli raise TH and IL-10 expression

CpG-ODN 1826 increase activation markers, TH and IL-10 expression

TH inhibitor has no effect on B cell survival

PD-L1/FasL-expression is not influenced by catecholamines

Activation with anti-IgM/CpG increase TH and PNMT expression in B cells

## Acknowledgements

I thank Katharina Krebber for designing primers for qRT-PCR.

## Author contributions

N.H. wrote the first draft of the manuscript, analyzed the data, designed the figures, established and performed experiments, G.P. and N.H., conceived the study, planned experiments, discussed results and revised the manuscript. B.O: established and performed experiments, discussed results and revised the manuscript. A.L: performed one experiment, discussed results and revised the manuscript. N.S., J.R.T, T.L., M.S. discussed results and revised the manuscript.

## Competing interests

The authors have declared that no competing interests exist.

## Data availability

All data supporting the findings of this study are available from the corresponding authors upon request.

## Funding

N.H. was supported by the Deutsche Forschungsgemeinschaft (DFG) fellowship (PO801/8-1). The funders had no role in study design, data collection and analysis, decision to publish, or preparation of the manuscript.

## Supplementary Information

**Supplementary Figure 1. Increased concentrations of TD-/TI-stimuli raise B cell activation**

(a, b): B cells were activated with different concentrations of the T cell-dependent (TD) mitogen (a: anti-CD40/IL-4) or T cell-independent (TI) mitogens (b: Poly I:C, TLR3; LPS TLR4) for 24h. As control group non-activated B cells were used. The expression of B cell activation markers MHC-II, CD86 and CD40 were determined on the surface of B cells by FACS (a; n = 5 and b; n = 4). For the experiments B cells from naïve DBA/1J mice were used. One-way analysis of variance (ANOVA) was used for comparisons. n.s. not significant; *p <0.5; **p < 0.01; ***p < 0.001; ****p < 0.0001.

**Supplementary Figure 2. Increased concentrations of TD-/TI-stimuli raise TH and IL-10 expression**

(a, b): B cells were activated with different concentrations of the TD mitogen anti-CD40/IL-4 or different concentrations of the TI mitogens TLR3 (Poly I:C) or TLR4 (LPS) for 24h. As control group non-activated B cells were used. (a): The frequency of CD19+TH+ B cells was measured by flow cytometry (TD: n = 6; TI: n = 4) and (b) the production of IL-10 was analyzed by ELISA (TD: n = 6; TI: n = 4-6). For the experiments B cells from naïve DBA/1J mice were used. One-way analysis of variance (ANOVA) (a, b) was used for comparisons. n.s. not significant, *p <0.5; **p < 0.01; ***p < 0.001; ****p < 0.0001.

**Supplementary Figure 3. CpG-ODN 1826 increase activation markers, TH and IL-10 expression**

(a-c): B cells were activated with different classes of CpG-Oligodesoxynucleotides (CpG-ODNs: ODN 1585 (class A); ODN 1826 (class B) and ODN 2395 (class C)) for 24h. Non-activated B cells treated with control ODNs (C-ODNs) were used as controls. The frequency of PI-CD19+MHC-II^high^+, PI-CD19+CD86+ and PI-CD19+CD40^high^+ B cells (a; n = 4) and CD19+TH+ B cells (b; n = 4) were analyzed by flow cytometry. The amount of IL-10 in cell culture supernatants was determined by ELISA (c; n = 8). For the experiments B cells from naïve DBA/1J mice were used. Data are pooled from four experiments (c). Student`s *t*-test (a-c) was used for comparisons. n.s. not significant, **p < 0.01; ****p < 0.0001.

**Supplementary Figure 4. TH inhibitor has no effect on B cell survival**

(a, b): B cells were treated for 30 min. with 0,5 mM of the tyrosine hydroxylase inhibitor 3-Iodo-L-tyrosine before activation with anti-IgM/CpG for 24h. As control group B cells were left untreated. (a): The frequency of living (PI-Annexin V-), early apoptotic (PI-Annexin V+), late apoptotic (PI^+^Annexin V^+^) and necrotic cells (PI+Annexin V-) was analyzed with the Annexin V-FITC binding assay by flow cytometry (n = 6). One of six representative dot plots is shown. (b): The frequency of living B cells was determined by FACS (n = 6). For the experiments B cells from naïve DBA/1J mice were used. Student`s *t*-test (b) was used for comparisons. n.s. not significant.

**Supplementary Figure 5. PD-L1/FasL-expression is not influenced by catecholamines**

(a): DBA/1J mice were immunized with 100μl of an emulsion of collagen type II and complete Freund`s adjuvans. After 14-21 days, splenic B cells from immunized mice were isolated by magnetic activated cell sorting (MACS) and B cell-depleted splenocytes were used for co-culture experiments. B cells were pre-treated with norepinephrine (NE; 10μM), isoproterenol (Isopr.; 10μM) or left untreated and cultivated with CD3/CD28-activated splenocytes for 72h. The expression of PD-L1 and FasL was analyzed on co-cultured B cells by FACS (PD-L1: n = 12; FasL: n = 11). Data are pooled from four experiments. One-way analysis of variance (ANOVA) was used for comparisons. n.s. not significant.

**Supplementary Figure 6. Activation with anti-IgM/CpG increase TH and PNMT expression in B cells**

Full western blot images show TH (a), PNMT (b) and GAPDH (a, b) expression in non-activated (0h) and activated B cells (24h and 48h). One of three representative blots is shown (n = 3).

## Notes

### Competing Interest Statement

The authors have declared no competing interest.

