## Supplementary figures and images for "TLR9-activated B cells improve their regulatory function by endogenously produced catecholamines"

### Activation with anti-IgM/CpG increase TH and PNMT expression in B cells

#### Supplementary Figure 6

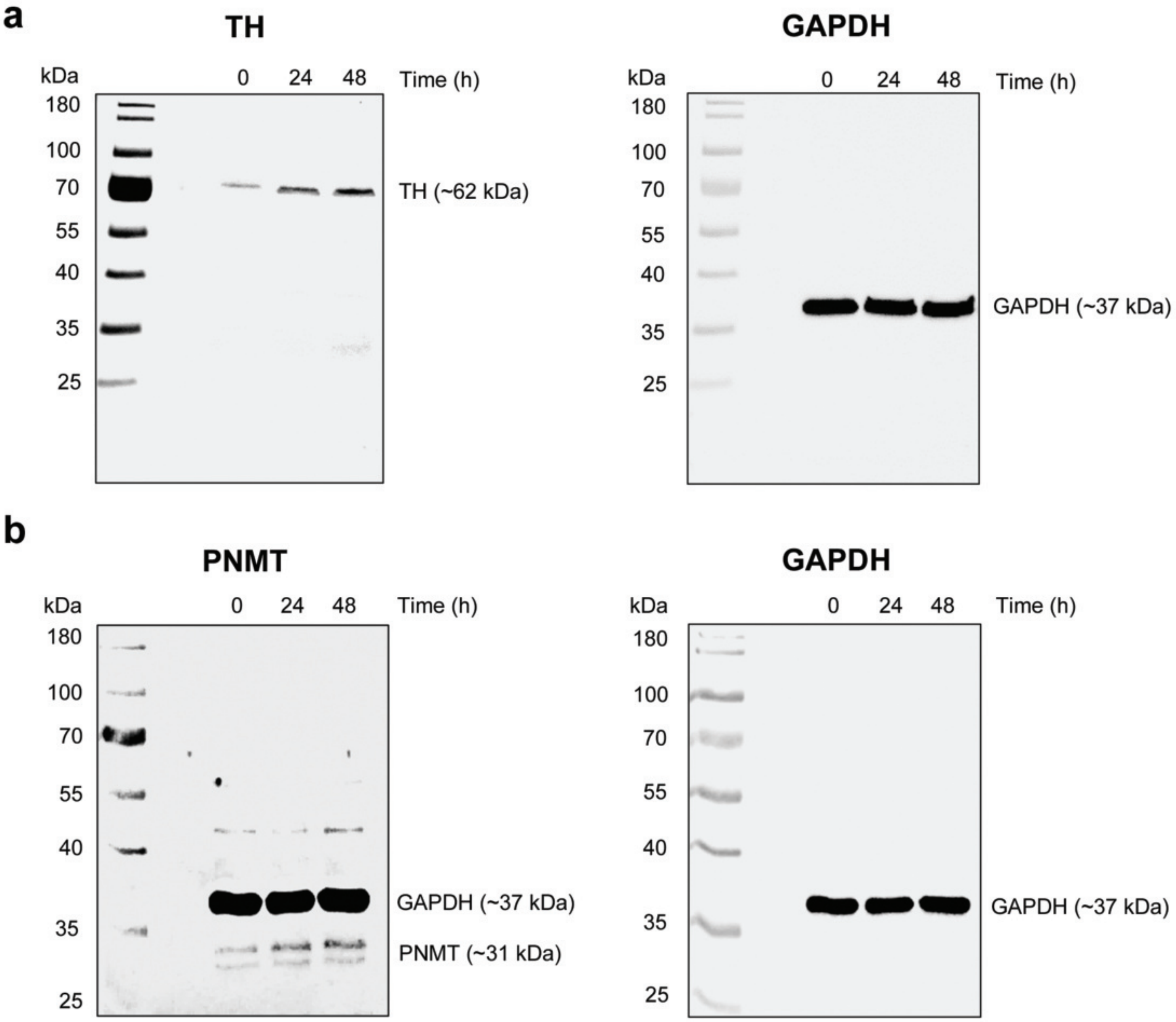

### CpG-ODN 1826 increase activation markers, TH and IL-10 expression

#### Supplementary Figure 3

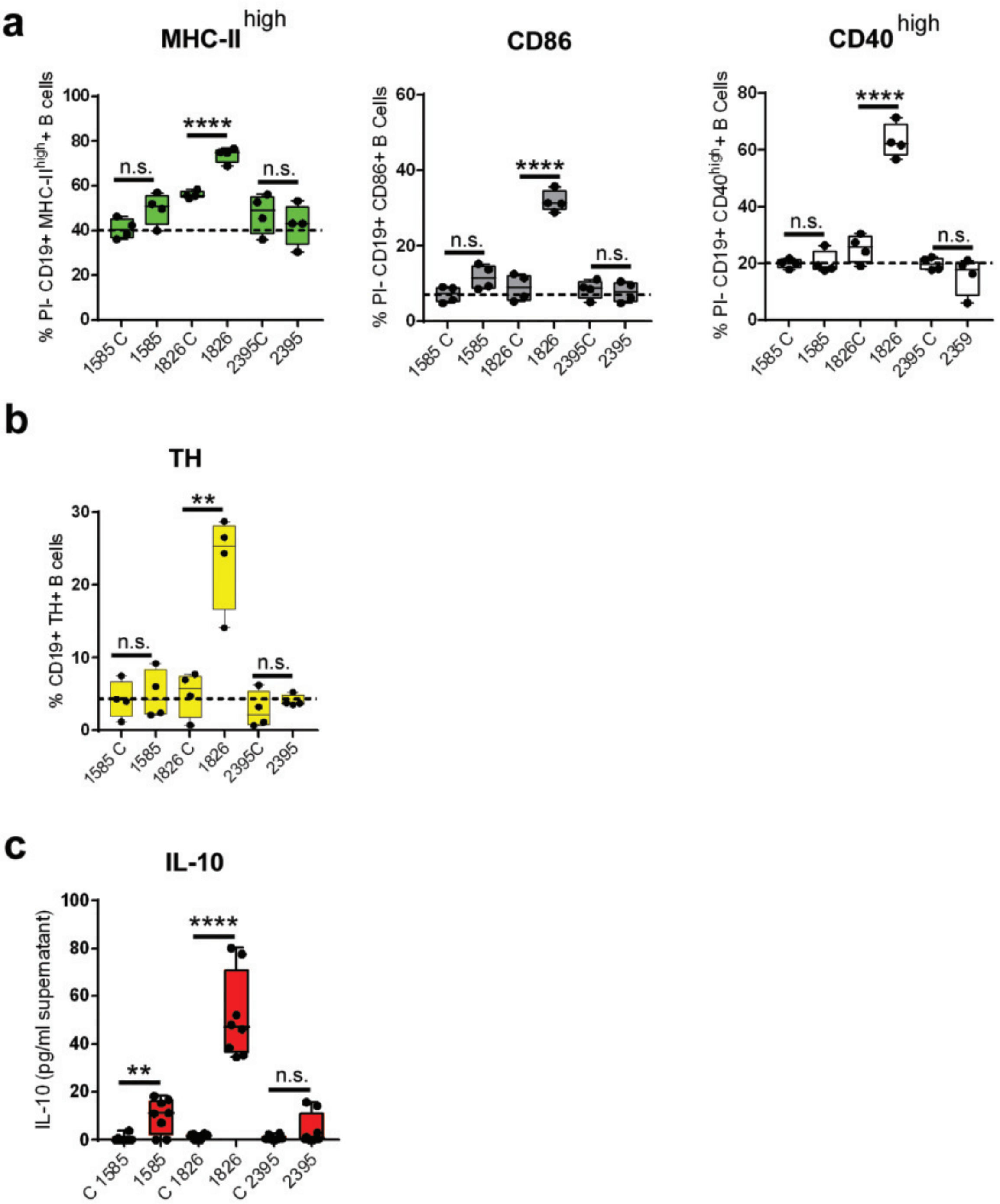

### Increased concentrations of TD-/TI-stimuli raise B cell activation

#### Supplementary Figure 1

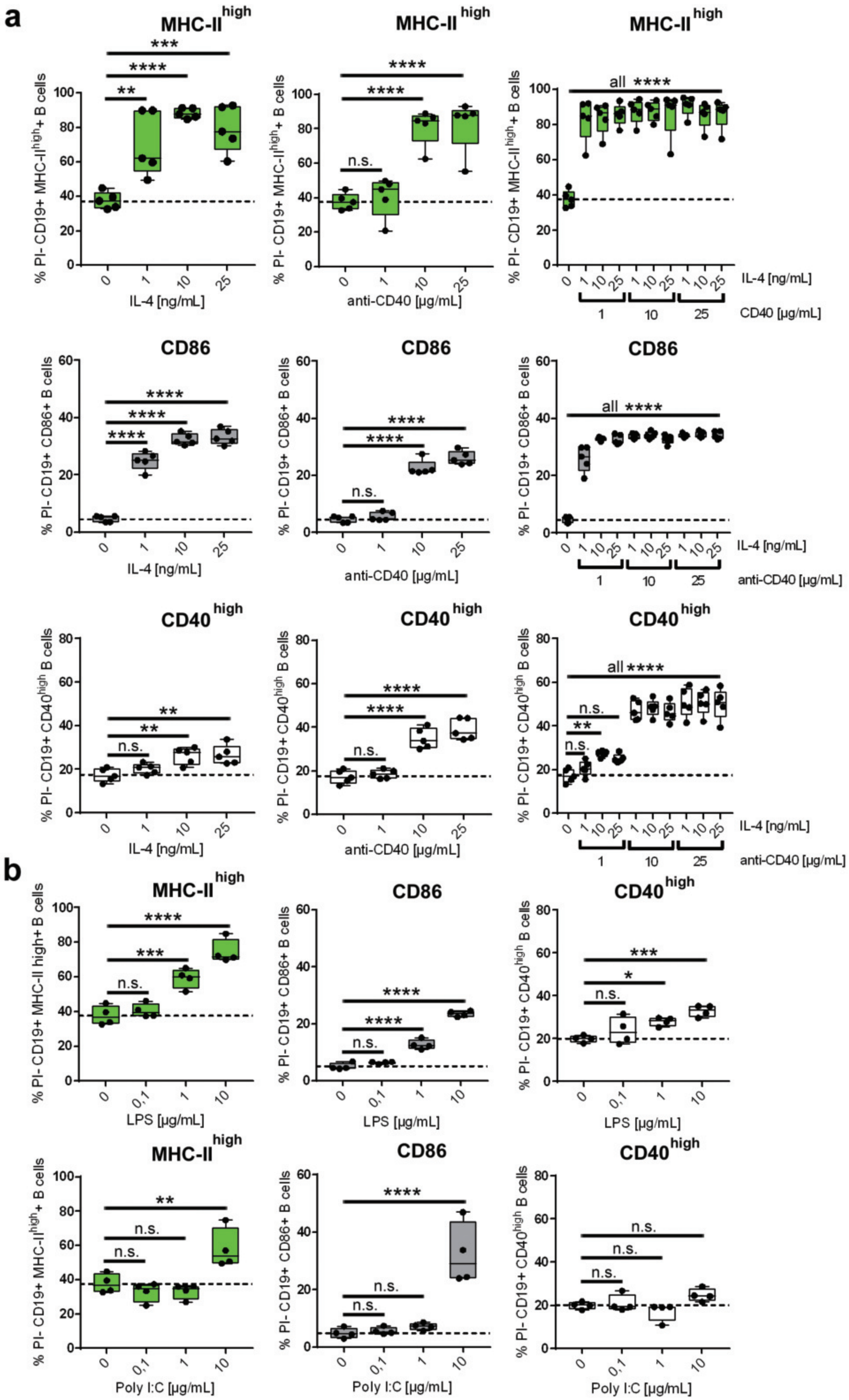

### Increased concentrations of TD-/TI-stimuli raise TH and IL-10 expression

#### Supplementary Figure 2

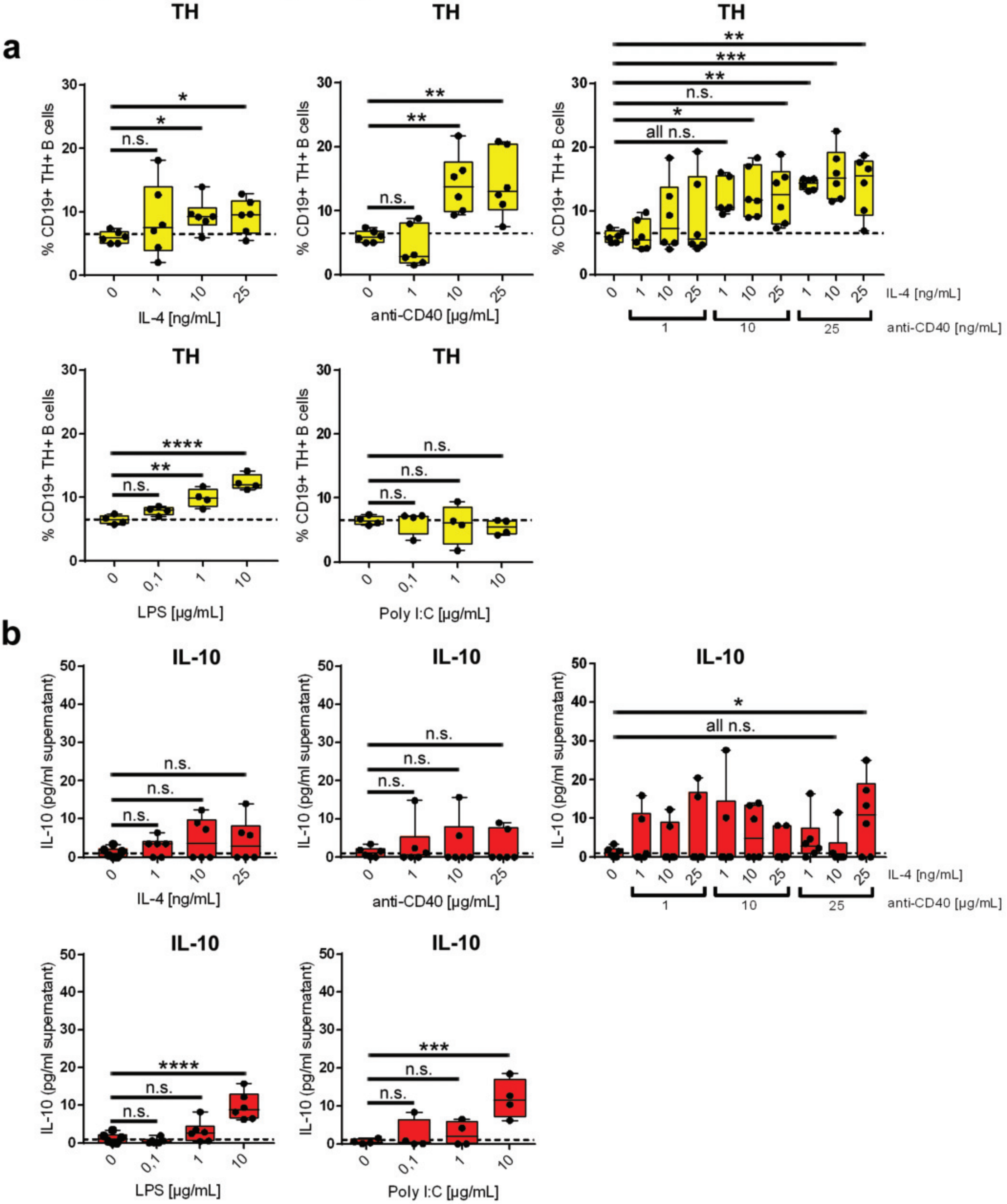

### PD-L1/FasL-expression is not influenced by catecholamines

#### Supplementary Figure 5

**a**

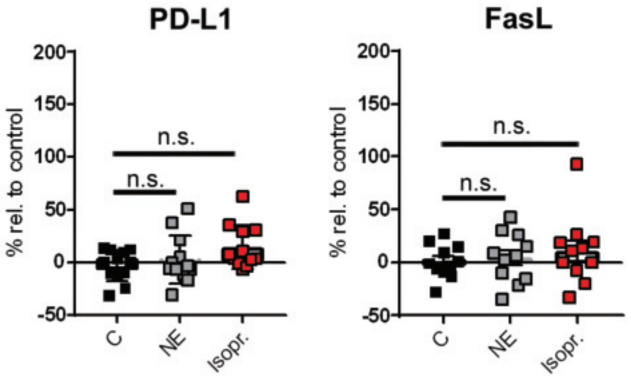

### TH inhibitor has no effect on B cell survival

#### Supplementary Figure 4

**a**

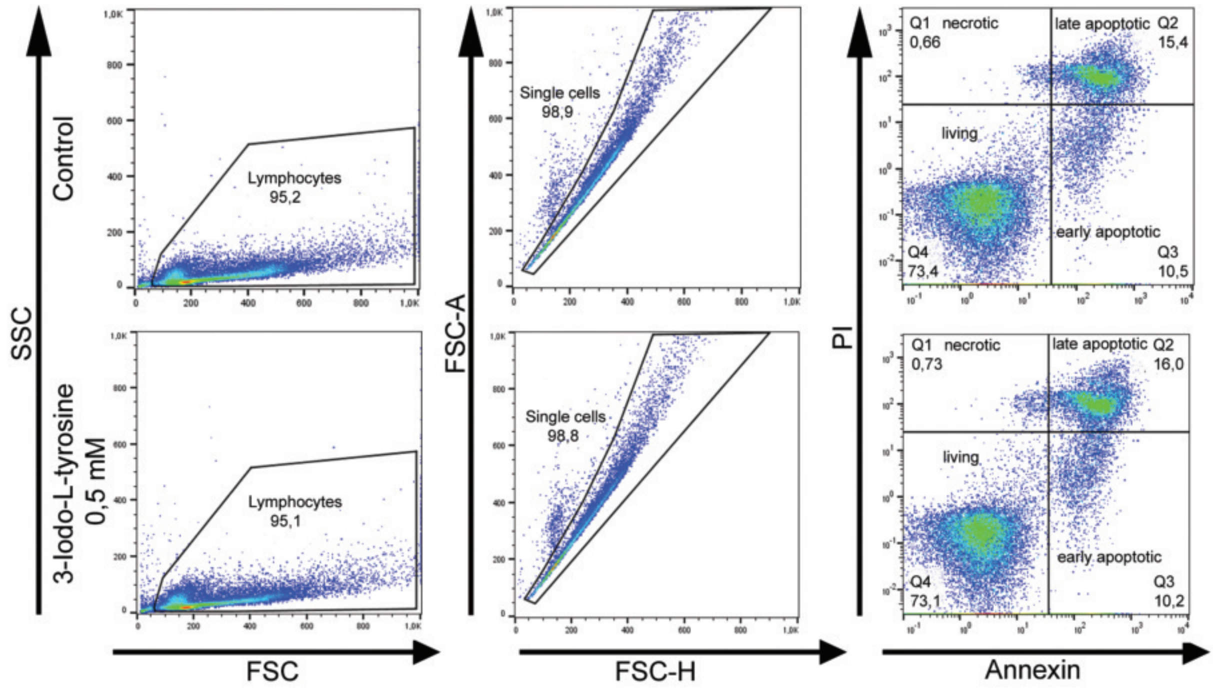


**b**

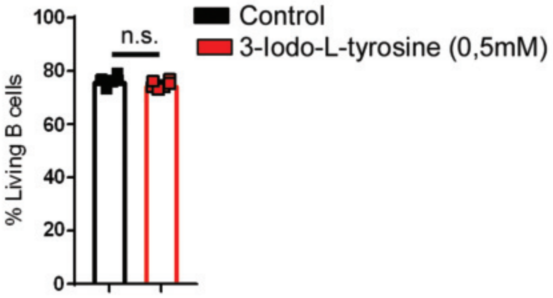
